# Generation and validation of an anti-human PANK3 mouse monoclonal antibody

**DOI:** 10.1101/2022.02.12.480208

**Authors:** Sunada Khadka, Long Vien, Paul Leonard, Laura Bover, Florian Muller

**Affiliations:** Department of Cancer Systems Imaging, The University of Texas MD Anderson Cancer Center, Houston, TX, USA; Department of Cancer Biology, The University of Texas MD Anderson Cancer Center, Houston, TX, USA; MD Anderson UT Health Graduate School of Biomedical Sciences, Houston, TX, USA; Department of Immunology, The University of Texas MD Anderson Cancer Center, Houston, TX, USA; Institute of Applied Cancer Science, The University of Texas MD Anderson Cancer Center, Houston, TX, USA; Department of Genomics Medicine, The University of Texas MD Anderson Cancer Center, Houston, TX, USA; Sporos Bioventures, Houston, TX, USA

**Keywords:** Pantothenate Kinases, PANK3, monoclonal antibody

## Abstract

Coenzyme A (CoA) is an essential co-factor at the intersection of diverse metabolic pathways. Cellular CoA biosynthesis is regulated at the first committed step— phosphorylation of pantothenic acid—catalyzed by pantothenate kinases (PANK1,2,3 in humans, PANK3 being the most highly expressed). Despite the critical importance of CoA in metabolism, the differential roles of PANK isoforms remain poorly understood. Our investigations of PANK proteins as precision oncology collateral lethality targets (*PANK1* is co-deleted as part of the *PTEN* locus in some highly aggressive cancers) were severely hindered by a dearth of commercial antibodies that can reliably detect endogenous PANK3. While we successfully validated commercial antibodies for PANK1 and PANK2 using CRISPR knockout cell lines, we found no commercial antibody that could detect endogenous PANK3. We therefore set out to generate a mouse monoclonal antibody against human PANK3 protein. We demonstrate that clone (Clone MDA-299-62A) can reliably detect endogenous PANK3 protein in cancer cell lines, with band-specificity confirmed by CRISPR PANK3 knockout cell lines. Sub-cellular fractionation indicates that PANK3 is overwhelmingly cytosolic and expressed broadly across cancer cell lines. PANK3 monoclonal antibody MDA-299-62A should prove a valuable tool for researchers investigating this understudied family of metabolic enzymes in health and disease.

## Introduction

Coenzyme A (CoA) is a central co-factor for more than 4% of known cellular enzymes and is involved in hundreds of biochemical reactions in the mammalian cells (1). As an important acyl group carrier, CoA is indispensable for reactions of the central metabolic pathways such as synthesis and oxidation of fatty acids, oxidative anaplerosis of pyruvate in the TCA cycle and the synthesis of lipids, isoprenoids and sterols, amino acid metabolism, and porphyrin synthesis (1,2). Additionally, CoA derived phosphopantetheine prosthetic group is also critical for enzymes involved in fatty acid, nonribosomal peptide, and polyketide synthesis(1,3,4). Pantothenate kinases control the first and the committed step of de novo CoA biosynthesis (5). Being the rate-limiting enzymes of the pathway, PANKs phosphorylate pantothenic acid (vitamin B5), which is converted to CoA in a series of four evolutionarily conserved enzymatic reactions (1,5). The family of PANKs constitutes four catalytically active isoforms—nuclear PANK1α and cytosolic PANK1β (both encoded by *PANK1*), mitochondrial (intermembrane space) PANK2 and cytosolic PANK3 (6). PANK isoforms are also differentially expressed and regulated, enabling these proteins to sense and maintain the levels of CoA and its thioesters differentially in specific cellular compartment (2,7). PANK proteins have been studied extensively in the context of normal mammalian physiology and pathologies that arise due to their dysregulation (8-12). Inactivating mutations in PANK2 protein have been linked to a neurodegenerative condition called pantothenate kinase-associated neurodegeneration (PKAN) in humans (13-17). Similarly, cancers with homozygous (bi-allelic) deletion of the tumor suppressor *PTEN*, that are typically highly aggressive, poor prognosis and refractory to treatment (18-20), can show co-deletion of the neighboring gene *PANK1*, making the redundant PANK proteins attractive therapeutic targets in these cancers (21,22). Despite the obvious importance of PANKs in CoA biosynthesis, the lack of reliable antibodies that can detect and distinguish endogenous PANK isoform proteins, has significantly thwarted efforts to better understand their role in both normal physiology and in pathological contexts, as well as characterize them as therapeutic targets. Generation of antibodies against PANK proteins, especially PANK3, has been significantly challenging given the extreme homology between the typical host species such as mice/rabbits, and humans. Additionally, the structural similarities in the PANK isozymes pose additional challenges in generating isozyme-specific antibodies. While many commercial antibodies may detect unphysiological levels of overexpressed PANK proteins, few antibodies are able to detect endogenously expressed PANK. Through an extensive and laborious screening of commercially available antibodies, we validated the antibodies for PANK1 and PANK2 for the detection of endogenous proteins using PANK-specific CRISPR KO cell lines. However, we could not find commercial antibodies capable of detecting endogenous human PANK3 protein.

Here, we describe the generation, optimization and validation of a mouse monoclonal antibody (mAb) (MDA-299-62A) against human PANK3 protein. We demonstrate that MDA-299-62A can reliably detect purified recombinant human PANK3, and endogenous PANK3 protein in cancer cell lysates with band specificity validated by PANK3 CRISPR knockout cancer cell lines. The PANK3 antibody generated in this study could serve as a reliable tool for quantifying endogenous PANK3 protein and should prove to be valuable for studies investigating these enzymes in health and disease.

## Methods

### Protein Expression and Purification

The pET28a plasmid vector containing DNA sequences encoding the PANK3 protein (residues pro12 to Asn368) was purchased from Addgene (25518) and transformed into both *E. coli* BL21 (DE3) and *E. coli* Rosetta2 (DE3) strains. The *E. coli* cells were grown in Terrific Broth media at 37°C until the optical density at 600 nm reached 0.5 absorbance units. The temperature of the culture was then reduced to 18°C prior to induction of the recombinant protein expression by the addition of 1 mM isopropyl ß-D-1-thiogalactopyranoside (IPTG). All protein purification steps were performed at 4°C After harvesting the cells by centrifugation, the *E. coli* cells were lysed by re-suspending the pellet in 10 mM Tris pH 7.5, 0.5 M NaCl, 5% glycerol, 5 mM Imidazole, complete protease inhibitors (Roche), 30 µg mL^-1^ DNase and 500 µg mL^-1^ lysozyme at pH 7.5 and lysed by sonication. The lysate was centrifuged at 20,000 rpm for 45 min to remove insoluble material. The recombinant protein was purified from the clarified lysate using a 2 mL Ni-NTA column pre-equilibrated in 10 mM Tris pH 7.5, 0.5 M NaCl, 5% glycerol, 5 mM Imidazole buffer. The Ni-NTA column was washed with 10 mM Tris-HCl, 0.5 M NaCl, 5% glycerol and 30 mM imidazole at pH 7.5. The bound proteins were eluted at 10 mM Tris-HCl, 0.5 M NaCl, 5% glycerol, 250 mM Imidazole at pH 7.5. The fractions were protein eluted were assessed by SDS PAGE.

### Monoclonal antibody production and purification

All animal experiments were performed using protocols approved by the Institutional Animal Care and Use Committee (IACUC). Antibody production using hybridoma technology and purifications were done at the MD Anderson Monoclonal Antibody Core. A detailed protocol on the protocol has been described. (23). Briefly, purified recombinant PANK3 protein was emulsified using the incomplete Freund Adjuvant (IFA, In-vivoGen) in a 1:1 ratio. Two NZBWF1/J mice and one BALB/c mice were used for immunization. 10 µg of recombinant protein (20 µl volume total) was administered through the foot-pad route every two days for the first five injections, and the dose was raised to 15 µg for three booster injections administered weekly. After completion of immunization, on day 31, mice were sacrificed and the popliteal lymph nodes and spleen from the immunized mice with the highest serum titer were harvested using sterile methods. B cell isolated either from the lymph node or the spleen were fused with Sp20 murine myeloma cells, and selected in hypoxanthine-aminopterin-thymidine (HAT) medium, and single cell clones were identified. Enzyme linked immunoassay (ELISA) screening was performed using the supernatant from each positive hybridoma clone to determine the efficiency of target protein binding by the mAb containing supernatants. Western blot was performed on HeLa cells to identify the clones that could detect endogenous PANK3 protein. The selected hybridoma clones were expanded and 300 mL of mAb rich supernatants were collected and filtered using a 0.45 µm membrane. The supernatants were added to Protein A columns for affinity chromatography purification. The antibodies captured in the column were eluted using elution buffer (0.1 M glycine-HCl, pH=3) that was neutralized with 1M Tris-HCl buffer. Overnight dialysis was performed in PBS to concentrate and preserve the antibody activity in a neutral buffer.

### ELISA Assay

ELISA plates were first coated with 100 µl of coating buffer containing the recombinant target protein used for immunization at 1 µg/ml and incubated at 4°C overnight. After 3X PBST wash, the plate was blocked with 100 µl of ELISA blocking buffer at RT for 1 hour. 50 µl of diluted serum from the immunized mice or the media supernatant from each hybridoma clone, which serve as the primary antibody, were added to individual wells respectively in the corresponding screenings, and incubated for 1 hour. The plate was then washed 3X in PBST, and 50 µl of goat anti-mouse HRP conjugated secondary antibody was added to each well and incubated for one hour. After 4X PBST washes, horseradish peroxidase (HRP) substrate TMB was added and the reaction was stopped by addition of H_2_SO4 after 30 mins. The OD 450 nm values were then measured using a spectrophotometer.

### Cell culture

The cell lines used in this work were HeLa (CVCL_0030, Cervical Carcinoma), 537 MEL (CVCL_8052, Melanoma), HAP1(CVCL_YO91), SKMEL-5 (CVCL_0527, Melanoma), SKMEL28 (CVCL_0526, Melanoma), SKMEL2 (CVCL_0069, Melanoma), G59 (CVCL_N729, Glioblastoma), D423-MG (CVCL_1160, Glioblastoma), referred to as D423 in the figures, LN319, a sub-clone of LN-99267, (CVCL_3958, Glioblastoma), D502 (CVCL_1162, Glioblastoma) and HEK293T (CVCL_0063). The cell lines were authenticated at the MD Anderson Cytogenetics and Cell Authentication Core. The cell lines were obtained from the following sources: HeLa (ATCC), 537 MEL (NCI), HAP1 (Horizon Discovery), SKMEL-5 (ATCC), SKMEL-2 (ATCC), SKMEL-28 (ATCC), G59 (Prof.Dr. Katrin Lamszus, Universitätsklinikum Hamburg-Eppendorf), D423 (Darrel Bigner, Duke University), LN319 (ATCC), D502 (ATCC), HEK 293T(ATCC). All cells were maintained in RPMI medium with 2mM glutamine (Cellgro/Corning #10-040-CV) supplemented with 10%FBS (Gibco/Life Technologies #16140-071) and 1% pen-strep (Gibco/Life Technologies#15140-122).

### Generation of CRISPR KO Clones

CRISPR KO was performed using the Santa Cruz dual plasmid CRISPR system. PANK CRISPR plasmids (PANK1 (sc-408890), PANK2 (sc-405120) and PANK3 sc-409325) and PANK HDR plasmids (PANK1 (sc-408890-HDR), PANK2 (sc-405120-HDR) and PANK3 (sc-409325-HDR) plasmids were purchased from Santa Cruz Biotechnologies. 1 µg of PANK plasmids and 1µg of PANK HDR plasmids were mixed together in 140 µl plasmid transfection medium (sc-108062). In a separate tube, 5 µl of UltraCruz® (sc-395739) transfection reagent was added to 145 µl transfection medium, and the plasmid and transfection reagents were mixed together and incubated for 10 minutes at RT. The cell culture medium in the target cells was replaced with fresh medium and the CRISPR/HDR plasmid mix were slowly added to the plates, and the plates were gently shaken to allow for a homogenous mixing. 24 hours later the medium was replaced with fresh medium and the cells were subjected to puromycin selection for a week. The cells were then trypsinized, washed in PBS and resuspended in FACS buffer (1%FBS+0.5mM EDTA in PBS). Single cell sorting was performed with RFP channel as a gate using a BD FACS ARIA Cells sorter. Confirmation of knockout was done with western blots.

### Generation of stable dox inducible PANK3 shRNA cell lines

Third generation lentivirus packaging system was used to generate the stable dox inducible shRNA expressing cells. Recombinant lentiviral particles were generated by transient transfection of HEK 293T cells following a standard protocol. ShRNA plasmids encoding a doxycycline inducible shRNA, hygromycin resistance marker and red fluorescent protein (RFP) were purchased from Cellecta. For transfection, 10 µg of the shRNA plasmid, were mixed with 5 µg of pMd2g, and 5 µg of pREV and 5 µg of p8.74 and 10 µg of shRNA vector and the plasmids were transfected into HEK 293T cells plated in 100 mm^2^ dishes. Fresh medium was replaced after 12 hours. Viral supernatants were collected, and fresh medium added on the HEK 293T cells every 24 hours until 48 hours since first media change. Cell debris were eliminated by centrifugation at 500g, and the supernatant containing the virus particles was sterile filtered through 0.45 µm filter. To infect the target cells, the filtered viral particles were mixed with an equal volume of fresh medium and polybrene (8 µg/ml) and added to target cells. The media on target cells were replaced with fresh media 24 hours after infection. After 48 hours since infection, 200 µg /ml hygromycin was added for selection. Knockdown of PANK3 was induced by addition of 2µg/ml dox for 48-72 hours and knockdown was confirmed by western blot.

### Western Blot

#### Whole cell lysate preparation

Cells were grown in 6 well plates for 48-72 hours and the lysates were harvested by washing the cells twice with ice cold phosphate-buffered saline (PBS). Ice-cold RIPA buffer with protease (cOmplete™ mini, Roche#11836153001) and phosphatase inhibitors (PhosSTOP, Roche, #5892970001), were added and the samples were then sonicated.

#### Cytosolic and mitochondrial Sub-cellular Fractionation

Sub-cellular fractionation was performed using the Thermo Fisher sub-cellular fractionation kit (#78840), using the manufacturer’s directions. Briefly, cells were trypsinized and the pellets were washed with ice-cold PBS. Ice cold cytoplasmic extraction buffer with 1X protease inhibitor cocktail was added to re-suspend the pellet and the sample was incubated on ice for 10 minutes with gentle mixing. The cell lysate was then centrifuged at 500g for 5 minutes at 4°C, and the cytosolic fraction in the supernatant was transferred to a pre-chilled tube, without agitating the pellet. Ice-cold membrane extraction buffer was added to the pellet, and the tube was vortexed at the highest setting for 5 seconds and incubated on ice for 10 minutes with gentle mixing. The tube was then centrifuged at 3000g for 5 minutes, and membrane fraction was isolated from the supernatant.

BCA assay (ThermoFisher, #23227), was used to determine the protein concentration. Proteins were separated by Nu-PAGE SDS-PAGE (4–12% gradient) and transferred onto nitrocellulose membranes using the semi-dry method (TransBlot turbo). Effective transfer of proteins on the membrane was verified with Ponceau S staining. 5% non-fat dry milk in tris-buffered saline (TBS) with 0.1% Tween 20 (TBST) was used as the blocking agent to block the non-specific sites on membrane. Primary antibodies (PANK1 CST 1:1000; PANK2 Origene 1:1000; PANK3 5µg/ml, TPI1 ProteinTech; 10713-1-AP 1:10000,) were added to the membranes and incubated overnight at 4°C with gentle rocking. On the second day, the membranes were washed 3x for 5 min with TBST. The membranes were then incubated in HRP tagged secondary antibody (1:5000) for 1 hour with gentle rocking and then washed 3x for 5 min with TBST. Then the membranes were incubated first with ECL substrate (32106) and X-ray films were exposed in a dark room and developed with different exposure times. If the bands were not visible, ThermoScientific SuperSignal West Femto (34096) was used as a substrate and the membranes developed on the X-ray films.

## Results

### 1. CRISPR-Validated commercial antibodies for PANK1 and PANK2, but not PANK3

To identify commercially available antibodies that can detect endogenous PANK proteins, we tested antibodies from diverse commercial suppliers. We found many commercial antibodies that can detect highly overexpressed unphysiological levels of PANKs, such as those obtained by transient transfection of PANK overexpressing plasmids. But very few can specifically detect endogenous expression of PANK. Additionally, the band specificities of these antibodies were not validated with a proper negative control by using a PANK deficient or PANK CRISPR knockout cell lines. To mitigate these issues, we purchased CRISPR knockout clones of PANK1, PANK2 and PANK3 in HAP1 cells (**Figure 1A**) from Horizon Discovery. HAP1 is a near-haploid cell line derived from KBM-7, a chronic myelogenous leukemia cell line. Due to its haploidy, HAP1 is an ideal model for genetic manipulations with CRISPR, and holds a great potential for genetic screening studies (24). Since HAP1 cells have almost one copy of most genes, the CRISPR induced heterozygous mutation artifacts (as in a diploid cell) are eliminated, allowing for easy and efficient generation of knockout clones. PANK isoform KOs in HAP1 cells were initially custom-made to order, but they are now available off-the shelf for other investigators to use. Additionally, we also independently generated knockout clones of PANK1, PANK2 and PANK3 in multiple different cancer cell lines to validate the band specificity and serve as a negative control for the antibodies (**Figure 1C-E**). We found that CST#23887S rabbit monoclonal antibody against PANK1 detects PANK1 protein at the expected size of 50 kD which disappears in two independent CRISPR knockout clones in HAP1 and HeLa cells (**Figure 1A** and **C**). Further validation of CST PANK1 antibody was also done in *PANK1* endogenous genomic deleted cell lines like 537 MEL and G59 and other *PANK1* intact cell lines, which further reinforced the band specificity of the antibody (**Figure 1B**). Similarly, we found that Origene’s #TA501321 mouse monoclonal antibody against PANK2 yields a band at the expected molecular weight (48 kDa), which disappears in the PANK2 CRISPR KO clones in HAP1, HeLa and LN319 cells (**Figure 1A, E** and **F**). However, we were unable to validate the loss of the bands of PANK1 and PANK2 proteins in the CRISPR KO clones by ProteinTech antibody (data not shown). We also tested multiple commercially available antibodies against PANK3 using PANK3 knockout CRISPR cell lines as negative controls. However, none of these antibodies gave the correct size band, and did not show loss of the band in CRISPR PANK3 KO cell lines. This led us to invest in generating our PANK3 antibody.

**Figure 1:**
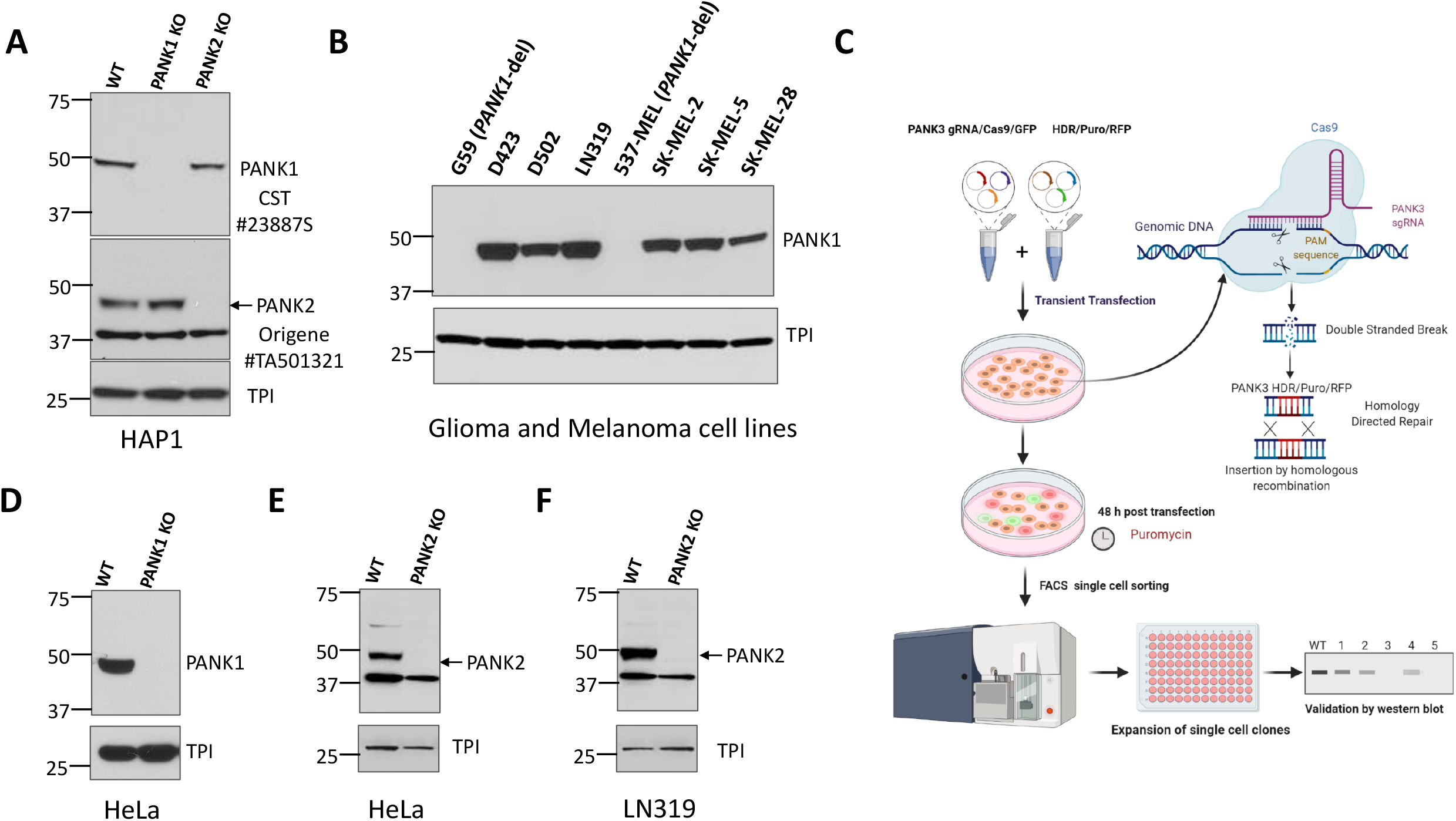
Validation of commercially available antibodies for PANK1 and PANK2. **A**. Commercially obtained and genomically-validated CRISPR knockout (KO) HAP1 cell lines were used as negative controls to verify specificity of PANK antibodies. Western blot confirming the loss of signal of PANK1 and loss of PANK2 in HAP1 PANK1 and PANK2 CRISPR KO clones purchased from Horizon Discovery. Rabbit anti-PANK1 mAb from CST (#23887S) correctly identifies a single band at 50 kDa (expected molecular weight of PANK1 protein) in WT HAP1 cells, and the band is absent in the PANK1 KO CRISPR clone. Origene mouse anti-PANK2 (mAb#TA501321) identifies endogenous PANK2 protein in wild type HAP1 cells at the expected size of 48 kD, and the band disappears in PANK2 CRISPR KO clone. A non-specific band at 37 kD is evident but is sufficiently distinct not to confound interpretation. Triose Phosphate Isomerase (TPI) is used as a loading control. **B**. Validation of CST PANK1 antibody using the PANK1 deleted and PANK1 intact glioma and melanoma lines. G59 (glioma) and 537 MEL(melanoma) are PANK1 homozygous deleted cancer cell lines. **C**. Schematic showing the Santa Cruz two-plasmid (gRNA/Cas9/GFP and HDR/Puromycin/RFP) system mediated CRISPR KO. **D-F**. Independent in-house generation of PANK1 and PANK2 CRISPR KO in cancer cells, and validation by western blot.

### 2. Production and Purification of His-tagged recombinant PANK3 protein

PANK3 is a 41 kD cytosolic protein which is ubiquitously expressed in all cell types. PANK3 protein is highly homologous between humans and rodents (homology with mice-99.1%, homology with rabbit-100%) (**Supplemental Figure S1**), which necessitated the use of the full length human PANK3 protein as the antigen, to allow for multiple and maximal antigenic as well as conformational epitope recognition in mice. The pET28a plasmid encoding six Histidine residues fused to 364 amino acid residues of the human PANK3 protein (residues pro12 to Asn368) was used as a vector for recombinant PANK3 production (**Figure 2A-C**). pET28a plasmid was transformed into *E. coli* and His-PANK3 synthesis was induced by the addition of IPTG into the growth medium. The clarified lysates from the *E. coli* cells were purified using a Ni-NTA column, and the eluted fractions were assessed for the presence of the recombinant PANK3 protein by Coomassie staining of the SDS-PAGE (**Figure 2A, C** and **D**). An unidentifiable band was observed around 75 kDa, so the Ni-NTA column elutes were further subjected to a size exclusion gel chromatography to obtain purified recombinant PANK3 protein (**Figure 2E**).

**Figure 2:**
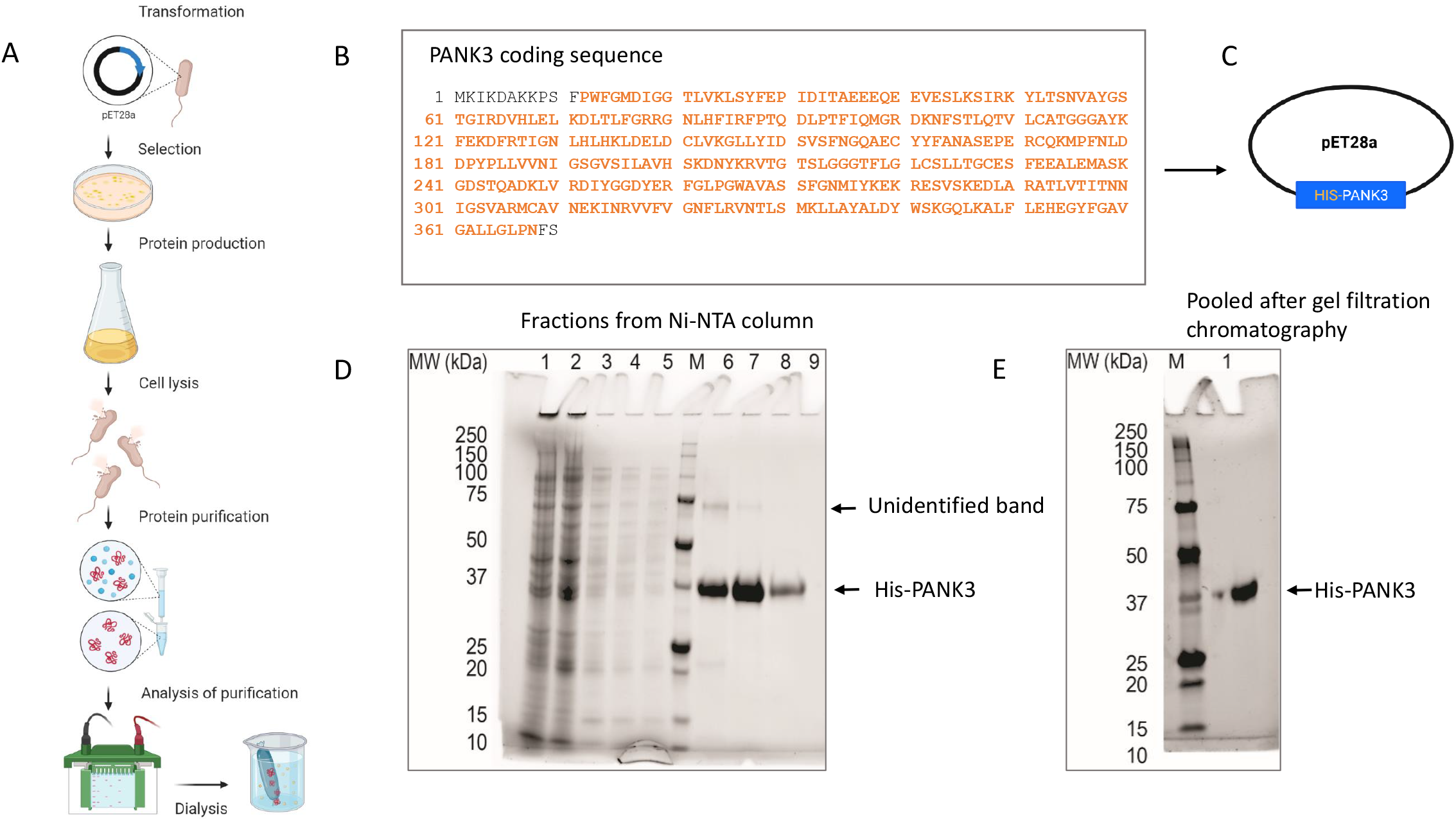
Generation and purification of recombinant PANK3 protein. **A-C**. Schematic showing the recombinant PANK3 production and purification. *E. coli* BL21 (DE3) and E. coli Rosetta2 (DE3) strains were transformed with pET28a plasmid vector encoding N terminal His tagged fusion PANK3 protein (residues pro12 to Asn368-highlighted in orange). The *E. coli* cells were grown in Terrific Broth media at 37°C overnight and then the temperature of the culture was reduced to 18°C prior to induction of the recombinant protein expression by the addition of 1 mM IPTG. Cells were harvested and lysed, and the soluble supernatant was applied in a pre-equilibrated Ni-NTA column, to allow the histidine tag to bind to the column. After washing the column, the bound proteins were eluted in 10 different fractions. **D-E**. SDS PAGE gel was stained with Coomassie Blue to assess the presence of His-PANK3 in the fractions. PANK3 protein purified from the Ni-NTA column was further subjected to gel filtration chromatography. Purified PANK3 protein eluted through gel filtration was validated by SDS-PAGE gel and coomassie staining.

### 3. Generation and validation of an anti-hPANK3 mouse mAb

To generate human PANK3 specific antibody, we immunized three mice—two NZB mice and a BALB/c mouse, with recombinant PANK3 protein (**Figure 3A**). Due to a 99.1% sequence homology in PANK3 protein between human and mouse (**Supplemental Figure S1**), we included the auto-immune NZB mouse model along with the wild type BALB/c mouse for immunization. The mice were administered 5 doses of PANK3 antigen mixed 1:1 with IFA every 2 days for the first 15 days, and then the final 3 booster doses were administered every week (**Figure 3A**) The serum samples collected from the mice were subjected to ELISA assay for quantification of serum PANK3 antibody titer. Due to high PANK3 antibody titer in NZB mouse #2, we generated hybridomas using the plasma cells isolated from this mouse (**Figure 3B-C**). We fused the splenocytes isolated from the immunized mouse with the murine myeloma cells Sp20 and selected the positive hybridoma clones in the HAT medium (**Figure 3C**).

**Figure 3:**
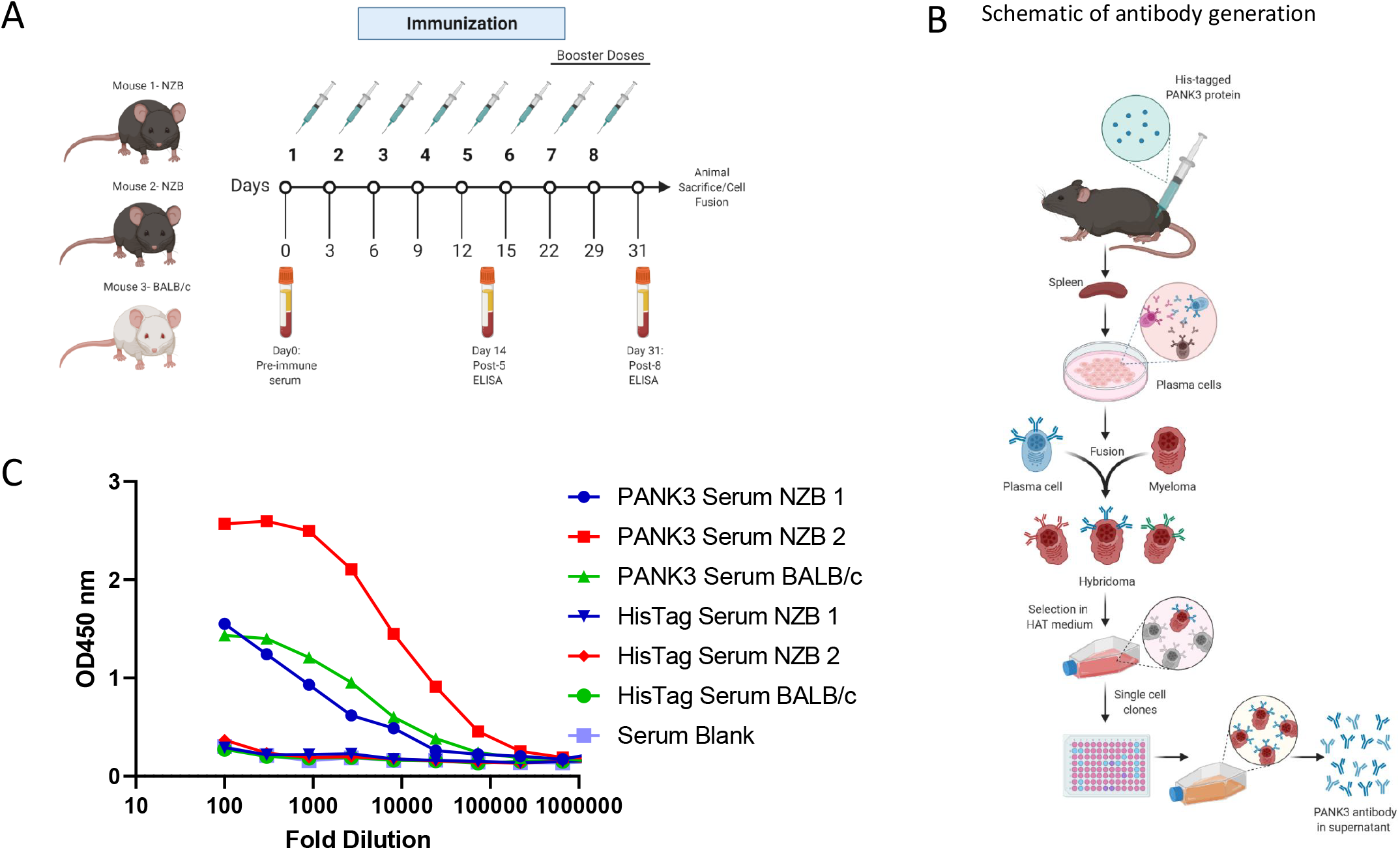
Generation of PANK3 antibody and detection of serum antibody levels by ELISA. **A**. Schematic showing the schedule of PANK3 protein immunization and blood collection for ELISA. PANK3 was administered in the foot pad of three mice, two NZB mice and a BALB/c mouse every two days for the first five doses and then weekly for three booster doses. Blood was collected on day 0, 14 and 31 to determine antibody concentration in the serum by ELISA. **B**. Schematic depicting the general workflow for the generation of PANK3 antibody. Briefly, recombinant 20 ul human PANK3 protein and IFA mix was injected into the foot pad of 3 mice (two NZB and one BALB/c). Following eight injections, mice were euthanized and plasma cells from the spleen were isolated and were fused with murine myeloma cells, Sp20 to form fusion hybridoma cells. After selection in HAT medium, the antibody producing plasma cells were single cell cloned and media supernatant from each clone was subjected to ELISA screen. **C**. Serum titration of anti-PANK3 antibody collected before mice sacrifice on day 31 using ELISA assay. Serum levels of PANK3 antibody was higher in mouse no.2, which was used for subsequent hybridoma generation, single clone isolation and antibody production. Serum response to His tag is compared to blank.

To determine the activity and efficacy of each hybridoma clone, we performed ELISA assay using the media supernatant from the hybridoma clones (**Figure 4A-B**). We further tested the efficacy and specificity of each clone-derived supernatant to detect endogenous PANK3 protein in HeLa cell lysates. Consistent with the ELISA results, the supernatants from most clones were able to recognize the recombinant antibody by immunoblotting assay (**Figure 4C**). In whole cell lysates, some of the clones were able to detect a band approximately at the expected molecular weight (∼41 kDa), however, none of the clones yielded clean and specific band for the endogenous PANK3 protein. We chose four clones that optimally recognized the endogenous PANK3 protein for further purification. As evidenced in **Figure 4C**, all four clones were able to detect the recombinant PANK3 protein, but the sub-optimal specificity for endogenous PANK3 and the presence of many non-specific bands posed significant challenges in further application of the antibody. To mitigate these issues, we ventured on optimizing the antibody and the immunoblotting assay conditions.

**Figure 4:**
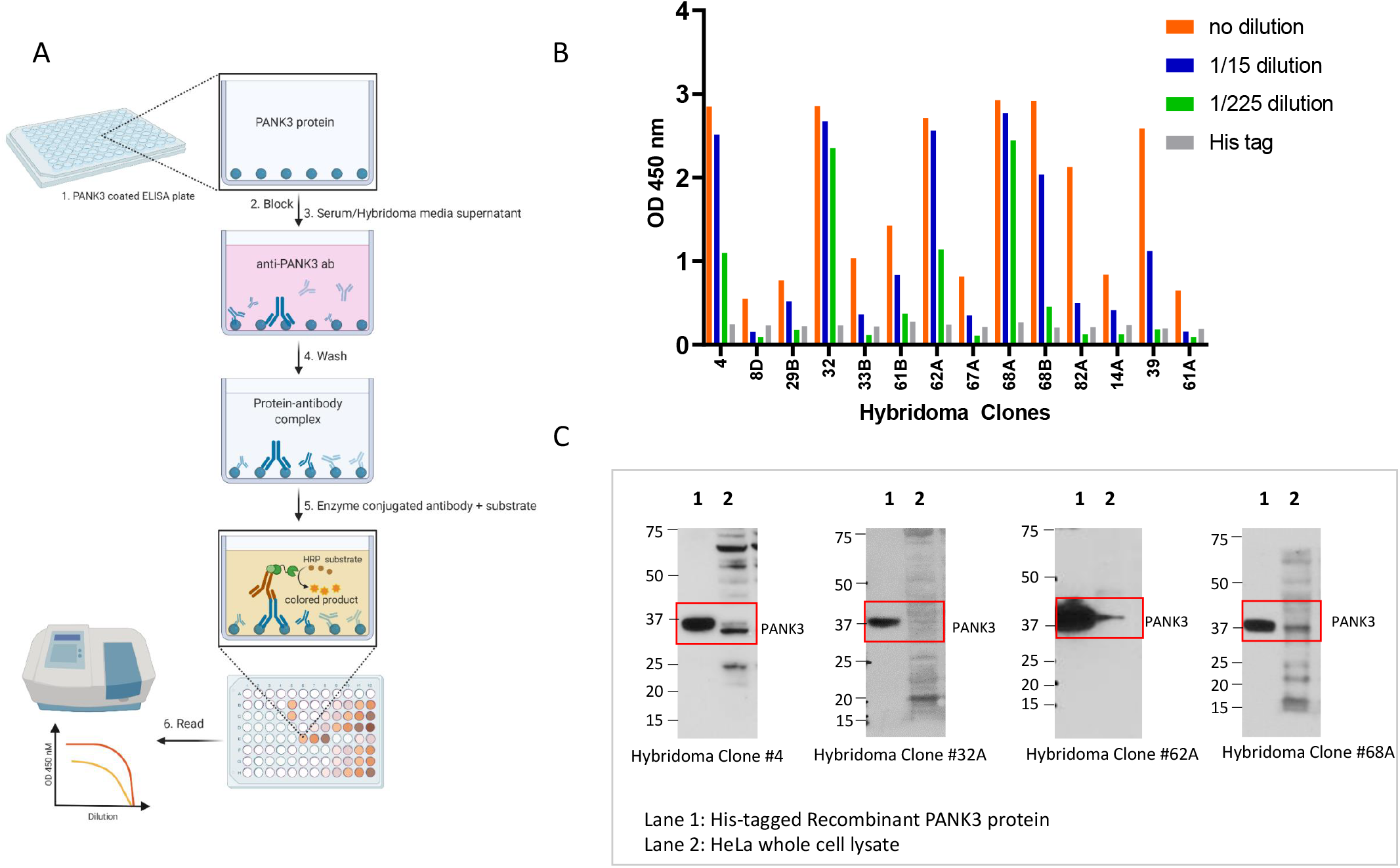
Quantification of PANK3 antibody levels in media supernatant of hybridoma clones. **A**. Schematic showing the detection of PANK3 protein by ELISA. Briefly, ELISA plates were coated with recombinant PANK3 protein and media supernatant (primary antibody) isolated from different hybridoma clones or the serum extracted from the immunized mice were added to the wells. An HRP linked secondary antibody was added to detect the PANK3 antibody bound to the immobilized PANK3 protein on the well. Addition of the substrate yielded a color change and the absorbance was quantified using a spectrophotometer. **B**. Absorbance values of medium supernatant (1:15 serial dilutions) isolated from different hybridoma clones. Only 14 clones out of 27 are shown in the graph. (See supplemental for all 27 clones). **C**. Immunoblot showing 4 antibody clones that were selected for further purification. The antibody clones are able to recognize recombinant protein, but the endogenous protein band in HeLa cell lysate is convoluted by overlapping non-specific bands.

### 4. Method-of-use optimization of MDA-299-62A anti-human PANK3 mouse mAb

PANK3 is a cytosolic protein in eukaryotes (**Figure 5A**). To improve the specificity of the antibody in immunoblotting assay, we performed subcellular fractionation to separate cytosolic protein including PANK3 from the total cell lysate. As shown in **Figure 5A-B**, we found that separating the cytosolic proteins by sub-cellular fractionation significantly reduced the non-specific signals and allowed reliable detection of endogenous PANK3 proteins in a broad range of cancer cell lines. Out of the four hybridoma clone supernatants that we purified, clone MDA-299-62A appeared to be the most effective in detecting endogenous PANK3 protein, with the most minimal signal to noise ratio. Therefore, we used this clone for further optimization. To conclusively validate that the antibody was indeed recognizing the endogenous PANK3 protein, we employed the CRISPR Cas9 technology to knock out PANK3 protein in two different cancer cell lines— HAP1 and HeLa (**Figure 5B and C**). We successfully obtained multiple single cell PANK3 knockout clones from these cell lines and validated that the antibody detected the correct band (**Figure 5B and C**). The CRISPR experiment also evidenced that PANK3 is a dispensable gene in most cancer cells, which is consistent with large-scale CRISPR and RNAi screen (DepMap) as well as mouse germline KO studies (2,25). We also performed knockdown studies using doxycycline inducible system and found that MDA-299-62A can reliably detect shRNA mediated knockdown of PANK3 in cancer cells.

**Figure 5:**
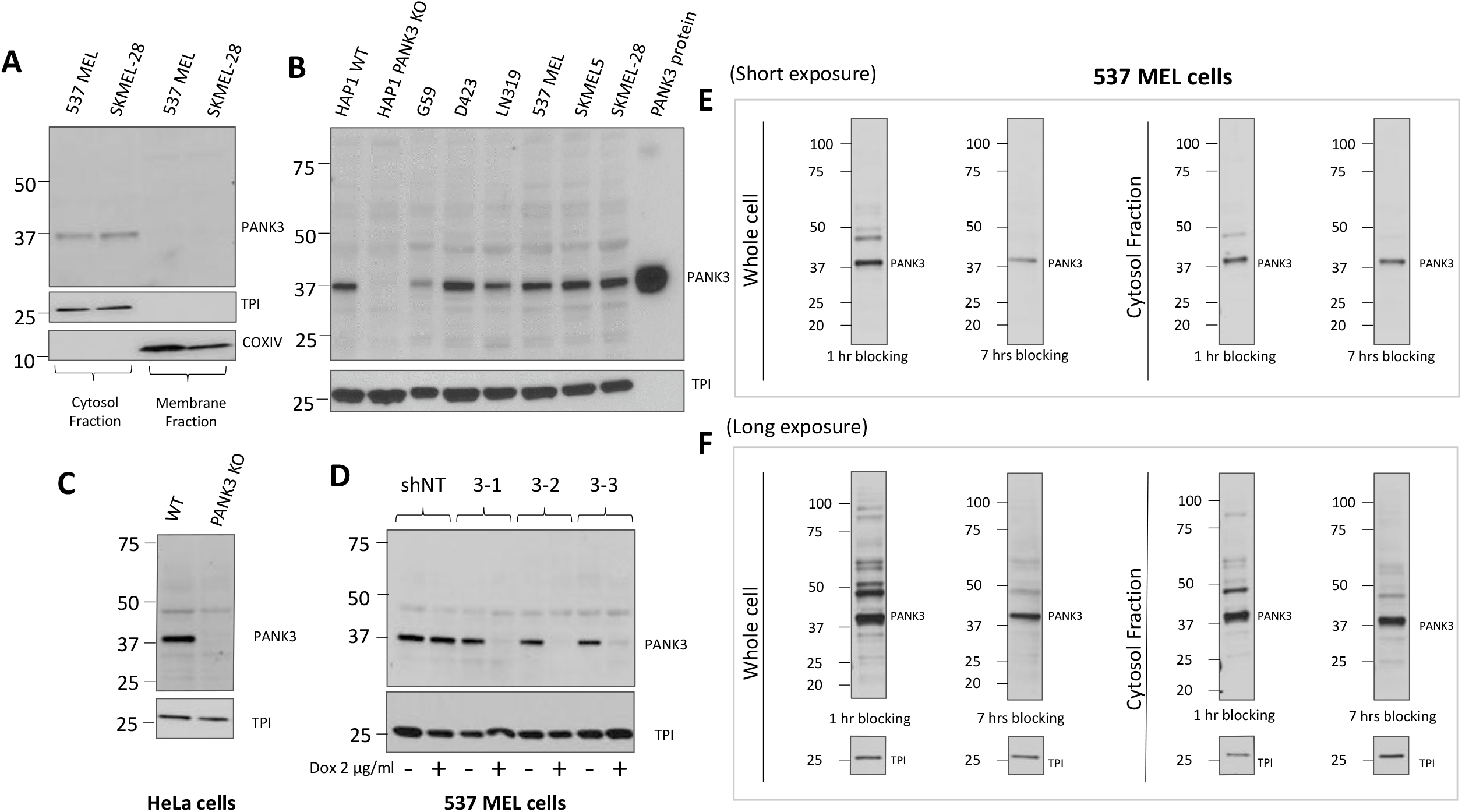
Optimization of PANK3 specific detection by subcellular fractionation of cells and longer blocking of transferred membranes. **A**. Immunoblots showing that PANK3 protein is found exclusively in the cytosolic fraction, confirming the cytosolic distribution of PANK3 protein in human cells. **B**. MDA-299-62A can detect the recombinant PANK3 protein (positive control) and endogenous PANK3 protein in cytosolic fractions of specified cancer cell lines. HAP1 PANK3 CRISPR KO(**B**) and HeLa PANK3 KO (**C**) cell lines were used as negative controls to verify correct band specificity. **D**. Knockdown of PANK3 protein by dox inducible shRNA, determined by immunoblot on cytosolic protein fractions using MDA-299-62A mouse monoclonal PANK3 antibody. **E-F**. The PANK3 band is detectable on whole cell lysates and the interference by non-specific bands is significantly reduced by increasing the duration of blocking to 7 hours instead of the typical 1 hour. Both whole cell lysates and cytosolic protein fractions are shown for reference, with shorter (15 secs) and longer exposure (5 minutes) of the x-ray film.

While subcellular fractionation could eliminate the non-specific signals on the immunoblot, it can be a cumbersome experiment to perform for routine use of PANK3 antibody for immunoblotting. Intending to reduce the non-specific signals without the need for subcellular fractionation, we modified the immunoblot assay protocol and employed a longer blocking duration of 6-8 hours, compared to the conventional 1 hour at RT on whole cell lysates (**Figure 5E and F**). We found that the duration of blocking could significantly dampen the non-specific signals and improved the signal to noise ratio in both cytosolic protein fractions as well as whole cell lysates (**Figure 5E and F**).

## Discussion

CoA is an indispensable cofactor for a myriad of biochemical reactions in the cells. Therefore, aberrations in its biosynthesis have been linked to debilitating neuropathology called PKAN in humans. Multiple therapeutic approaches are currently being explored in preclinical and clinical studies to restore CoA levels to combat CoA deficiency in these pathologies (14,15). PANK proteins control the first and the rate-limiting step in the de-novo coenzyme A biosynthesis. Owing to the essentiality of CoA at the cellular as well as organismal level, PANK and other CoA pathways proteins have also garnered interest as attractive therapeutic targets against parasites such as *Plasmodium falciparum* and *Toxoplasma gondii* (26). The importance of CoA in microbes and the distinctions in CoA biosynthesis pathway proteins between microbes and mammals have also propelled investigative interest on CoA biosynthesis as anti-bacterial targets (27,28). Our drug-target discovery investigations of PANK proteins also relies on the essentiality of PANK proteins for CoA biosynthesis and aims to exploit the redundancies in the PANK proteins to specifically target *PANK1*-deleted cancers, which can be co-deleted as part of the *PTEN* tumor suppressor locus, whose homozygous deletion is associated with poor prognosis, high malignancy and resistance to both conventional chemotherapy and precision oncology drugs (18-22). Despite significant investigative interest in understanding the roles of PANK proteins in normal and pathological conditions, the paucity of antibodies to detect endogenous PANK proteins has been a detriment to the advancement of these studies.

In this study, we report the validation of commercially available PANK1 and PANK2 antibodies and demonstrate detection of band specificity using PANK1 and PANK2 CRISPR KO cells. Since we were unable to obtain any reliable PANK3 antibody against endogenous human PANK3 protein, we generated and validated a human PANK3 specific mouse monoclonal antibody. We show that MDA-299-62A can reliably detect recombinant PANK3 protein and endogenous PANK3 protein in ELISA and immunoblotting assays. We confirmed the band specificity of our antibody by using a PANK3 CRISPR knockout cell line, Our results show that the antibody MDA-299-62A can be routinely used for immunoblotting assays, but with the caveat of multiple other unspecific bands along with the PANK3 protein in the cell lysate (though this is also the case with commercial anti-PANK2, mouse mAb (Origene #TA501321, **Figure 1E-F**). The use of full-length protein as an antigen possibly explains the detection of many unspecific bands by the antibody. PANK3 is extremely conserved between humans and rodents (rabbit 100%). So, the antibody was raised in mice, which shares a 99.1% homology to human PANK3 protein. We used the full-length protein in an autoimmune mouse model, to allow for maximum antigen recognition and antibody production. While developing the antibody in organisms with significant sequence differences could alleviate this issue, however, our attempt of developing PANK3 polyclonal antibody in chickens did not yield improved results (data not shown).

To alleviate the issue of unspecific bands recognition by MDA-299-62A, we demonstrate that by performing subcellular fractionation, we can concentrate PANK3 in the cytosol fraction, which can significantly minimize unspecific bands. Since performing sub-cellular fractionation on a routine basis can be cumbersome, we demonstrate that by extending the duration of blocking of the membranes, we can significantly minimize these unspecific bands. These mitigation measures are only relevant for immunoblotting assays, and further optimization of MDA-299-62A mAb will be required for immunohistochemistry, immunofluorescence as well as fluorescence activate cell sorting (FACS) experiments. However, the consistent and reliable results that we obtained with MDA-299-62A from well controlled experiments shows that the antibody is robust enough to study the genetic interactions of PANK isoform knockout for the purposes of validating PANK as collateral lethality targets in cancer.

## Author’s contributions

SK and FLM conceived the study. PL performed recombinant protein expression and purification. LV and LB performed animal immunization, hybridoma generation and antibody purification. SK performed all *in-vitro* genetic and antibody validation experiments. SK wrote the manuscripts with assistance from FLM. All authors approved the final manuscript.

## Data Availability

All data relevant to the manuscripts are included.

## Acknowledgements

We thank Dr. Ronald DePinho for his guidance and research support on this work. We also want to thank Sarah Joseph for her help in generation and purification of recombinant PANK3 protein. The graphical abstracts and figures were adapted from or created with Biorender.com.

## References

1. Leonardi, R., Zhang, Y. M., Rock, C. O., and Jackowski, S. (2005) Coenzyme A: back in action. Prog Lipid Res 44, 125–153

2. Dansie, L. E., Reeves, S., Miller, K., Zano, S. P., Frank, M., Pate, C., Wang, J., and Jackowski, S. (2014) Physiological roles of the pantothenate kinases. Biochem Soc Trans 42, 1033–1036

3. Gout, I. (2018) Coenzyme A, protein CoAlation and redox regulation in mammalian cells. Biochem Soc Trans 46, 721–728

4. Theodoulou, F. L., Sibon, O. C., Jackowski, S., and Gout, I. (2014) Coenzyme A and its derivatives: renaissance of a textbook classic. Biochem Soc Trans 42, 1025–1032

5. Jackowski, S., and Rock, C. O. (1981) Regulation of coenzyme A biosynthesis. J Bacteriol 148, 926–932

6. Alfonso-Pecchio, A., Garcia, M., Leonardi, R., and Jackowski, S. (2012) Compartmentalization of mammalian pantothenate kinases. PLoS One 7, e49509

7. Vallari, D. S., Jackowski, S., and Rock, C. O. (1987) Regulation of pantothenate kinase by coenzyme A and its thioesters. J Biol Chem 262, 2468–2471

8. Leonardi, R., Rehg, J. E., Rock, C. O., and Jackowski, S. (2010) Pantothenate kinase 1 is required to support the metabolic transition from the fed to the fasted state. PLoS One 5, e11107

9. Jackowski, S., and Leonardi, R. (2014) Deregulated coenzyme A, loss of metabolic flexibility and diabetes. Biochem Soc Trans 42, 1118–1122

10. Leonardi, R., Rock, C. O., and Jackowski, S. (2014) Pank1 deletion in leptin-deficient mice reduces hyperglycaemia and hyperinsulinaemia and modifies global metabolism without affecting insulin resistance. Diabetologia 57, 1466–1475

11. Corbin, D. R., Rehg, J. E., Shepherd, D. L., Stoilov, P., Percifield, R. J., Horner, L., Frase, S., Zhang, Y. M., Rock, C. O., Hollander, J. M., Jackowski, S., and Leonardi, R. (2017) Excess coenzyme A reduces skeletal muscle performance and strength in mice overexpressing human PANK2. Mol Genet Metab 120, 350–362

12. Subramanian, C., Yao, J., Frank, M. W., Rock, C. O., and Jackowski, S. (2020) A pantothenate kinase-deficient mouse model reveals a gene expression program associated with brain coenzyme a reduction. Biochim Biophys Acta Mol Basis Dis 1866, 165663

13. Sharma, L. K., Leonardi, R., Lin, W., Boyd, V. A., Goktug, A., Shelat, A. A., Chen, T., Jackowski, S., and Rock, C. O. (2015) A high-throughput screen reveals new small-molecule activators and inhibitors of pantothenate kinases. J Med Chem 58, 1563–1568

14. Zano, S. P., Pate, C., Frank, M., Rock, C. O., and Jackowski, S. (2015) Correction of a genetic deficiency in pantothenate kinase 1 using phosphopantothenate replacement therapy. Mol Genet Metab 116, 281–288

15. Sharma, L. K., Subramanian, C., Yun, M. K., Frank, M. W., White, S. W., Rock, C. O., Lee, R. E., and Jackowski, S. (2018) A therapeutic approach to pantothenate kinase associated neurodegeneration. Nat Commun 9, 4399

16. Hortnagel, K., Prokisch, H., and Meitinger, T. (2003) An isoform of hPANK2, deficient in pantothenate kinase-associated neurodegeneration, localizes to mitochondria. Hum Mol Genet 12, 321–327

17. Kotzbauer, P. T., Truax, A. C., Trojanowski, J. Q., and Lee, V. M. (2005) Altered neuronal mitochondrial coenzyme A synthesis in neurodegeneration with brain iron accumulation caused by abnormal processing, stability, and catalytic activity of mutant pantothenate kinase 2. J Neurosci 25, 689–698

18. Bucheit, A. D., Chen, G., Siroy, A., Tetzlaff, M., Broaddus, R., Milton, D., Fox, P., Bassett, R., Hwu, P., Gershenwald, J. E., Lazar, A. J., and Davies, M. A. (2014) Complete loss of PTEN protein expression correlates with shorter time to brain metastasis and survival in stage IIIB/C melanoma patients with BRAFV600 mutations. Clin Cancer Res 20, 5527–5536

19. Peng, W., Chen, J. Q., Liu, C., Malu, S., Creasy, C., Tetzlaff, M. T., Xu, C., McKenzie, J. A., Zhang, C., Liang, X., Williams, L. J., Deng, W., Chen, G., Mbofung, R., Lazar, A. J., Torres-Cabala, C. A., Cooper, Z. A., Chen, P. L., Tieu, T. N., Spranger, S., Yu, X., Bernatchez, C., Forget, M. A., Haymaker, C., Amaria, R., McQuade, J. L., Glitza, I. C., Cascone, T., Li, H. S., Kwong, L. N., Heffernan, T. P., Hu, J., Bassett, R. L. Jr.,, Bosenberg, M. W., Woodman, S. E., Overwijk, W. W., Lizee, G., Roszik, J., Gajewski, T. F., Wargo, J. A., Gershenwald, J. E., Radvanyi, L., Davies, M. A., and Hwu, P. (2016) Loss of PTEN Promotes Resistance to T Cell-Mediated Immunotherapy. Cancer Discov 6, 202–216

20. Catalanotti, F., Cheng, D. T., Shoushtari, A. N., Johnson, D. B., Panageas, K. S., Momtaz, P., Higham, C., Won, H. H., Harding, J. J., Merghoub, T., Rosen, N., Sosman, J. A., Berger, M. F., Chapman, P. B., and Solit, D. B. (2017) PTEN Loss-of-Function Alterations Are Associated With Intrinsic Resistance to BRAF Inhibitors in Metastatic Melanoma. JCO Precis Oncol 1

21. Muller, F. L., Aquilanti, E. A., and DePinho, R. A. (2015) Collateral Lethality: A new therapeutic strategy in oncology. Trends Cancer 1, 161–173

22. Muller, F. L., Colla, S., Aquilanti, E., Manzo, V. E., Genovese, G., Lee, J., Eisenson, D., Narurkar, R., Deng, P., Nezi, L., Lee, M. A., Hu, B., Hu, J., Sahin, E., Ong, D., Fletcher-Sananikone, E., Ho, D., Kwong, L., Brennan, C., Wang, Y. A., Chin, L., and DePinho, R. A. (2012) Passenger deletions generate therapeutic vulnerabilities in cancer. Nature 488, 337–342

23. Muhsin, A., Rangel, R., Vien, L., and Bover, L. (2022) Monoclonal Antibodies Generation: Updates and Protocols on Hybridoma Technology. Methods Mol Biol 2435, 73–93

24. Wutz, A. (2014) Haploid animal cells. Development 141, 1423–1426

25. Garcia, M., Leonardi, R., Zhang, Y. M., Rehg, J. E., and Jackowski, S. (2012) Germline deletion of pantothenate kinases 1 and 2 reveals the key roles for CoA in postnatal metabolism. PLoS One 7, e40871

26. Schalkwijk, J., Allman, E. L., Jansen, P. A. M., de Vries, L. E., Verhoef, J. M. J., Jackowski, S., Botman, P. N. M., Beuckens-Schortinghuis, C. A., Koolen, K. M. J., Bolscher, J. M., Vos, M. W., Miller, K., Reeves, S. A., Pett, H., Trevitt, G., Wittlin, S., Scheurer, C., Sax, S., Fischli, C., Angulo-Barturen, I., Jimenez-Diaz, M. B., Josling, G., Kooij, T. W. A., Bonnert, R., Campo, B., Blaauw, R. H., Rutjes, F., Sauerwein, R. W., Llinas, M., Hermkens, P. H. H., and Dechering, K. J. (2019) Antimalarial pantothenamide metabolites target acetyl-coenzyme A biosynthesis in Plasmodium falciparum. Sci Transl Med 11

27. Gerdes, S. Y., Scholle, M. D., D’Souza, M., Bernal, A., Baev, M. V., Farrell, M., Kurnasov, O. V., Daugherty, M. D., Mseeh, F., Polanuyer, B. M., Campbell, J. W., Anantha, S., Shatalin, K. Y., Chowdhury, S. A., Fonstein, M. Y., and Osterman, A. L. (2002) From genetic footprinting to antimicrobial drug targets: examples in cofactor biosynthetic pathways. J Bacteriol 184, 4555–4572

28. Butman, H. S., Kotze, T. J., Dowd, C. S., and Strauss, E. (2020) Vitamin in the Crosshairs: Targeting Pantothenate and Coenzyme A Biosynthesis for New Antituberculosis Agents. Front Cell Infect Microbiol 10, 605662

