## Supplemental Figures for "Generation and validation of an anti-human PANK3 mouse monoclonal antibody"

**A**

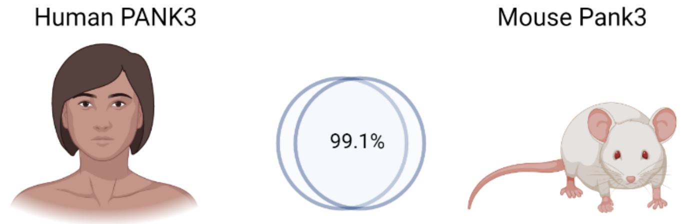


PANK3 1 MKIKDAKKPSFPWFGMDIGGTLVKLSYFEPIDITAEEEQEEVESLKSIRKYLTSNVAYGS 60
Pank3 1 MKIKDAKKPSFPWFGMDIGGTLVKLSYFEPIDITAEEEQEEVESLKSIRKYLTSNVAYGS 60

PANK3 61 TGIRDVHLELKDLTLFGRRGNLHFIRFPTQDLPTFIQMGRDKNFSTLQTVL**C**ATGGGAYK 120
Pank3 61 TGIRDVHLELKDLTLFGRRGNLHFIRFPTQDLPTFIQMGRDKNFSTLQTVL**S**ATGGGAYK 120

PANK3 121 FEKDFRTIGNLHLHKLDELDCLVKGLLYIDSVSFNGQAECYYFANASEPERCQKMPFNLD 180
Pank3 121 FEKDFRTIGNLHLHKLDELDCLVKGLLYIDSVSFNGQAECYYFANASEPERCQKMPFNLD 180

PANK3 181 DPYPLLVVNIGSGVSILAVHSKDNYKRVTGTSLGGGTFLGLCSLLTGCESFEEALEMASK 240
Pank3 181 DPYPLLVVNIGSGVSILAVHSKDNYKRVTGTSLGGGTFLGLCSLLTGCESFEEALEMASK 240

PANK3 241 GDSTQAD**K**LVRDIYGGDYERFGLPGWAVASSFGNMIYKEKRE**S**VSKEDLARATLVTITNN 300
Pank3 241 GDSTQAD**R**LVRDIYGGDYERFGLPGWAVASSFGNMIYKEKRE**T**VSKEDLARATLVTITNN 300

PANK3 301 IGSVARMCAVNEKINRVVFVGNFLRVNTLSMKLLAYALDYWSKGQLKALFLEHEGYFGAV 360
Pank3 301 IGSVARMCAVNEKINRVVFVGNFLRVNTLSMKLLAYALDYWSKGQLKALFLEHEGYFGAV 360

PANK3 361 GALLGLPNFS 370
Pank3 361 GALLGLPNFS 370

**B**

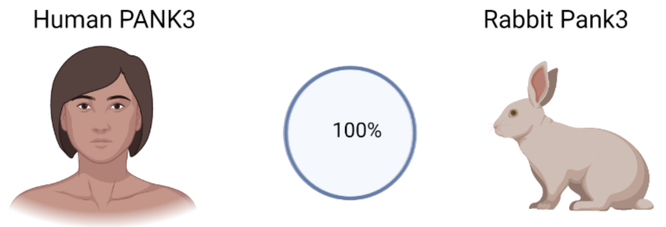


PANK3 1 MKIKDAKKPSFPWFGMDIGGTLVKLSYFEPIDITAEEEQEEVESLKSIRKYLTSNVAYGS 60
Pank3 1 MKIKDAKKPSFPWFGMDIGGTLVKLSYFEPIDITAEEEQEEVESLKSIRKYLTSNVAYGS 60

PANK3 61 TGIRDVHLELKDLTLFGRRGNLHFIRFPTQDLPTFIQMGRDKNFSTLQTVLCATGGGAYK 120
Pank3 61 TGIRDVHLELKDLTLFGRRGNLHFIRFPTQDLPTFIQMGRDKNFSTLQTVLCATGGGAYK 120

PANK3 121 FEKDFRTIGNLHLHKLDELDCLVKGLLYIDSVSFNGQAECYYFANASEPERCQKMPFNLD 180
Pank3 121 FEKDFRTIGNLHLHKLDELDCLVKGLLYIDSVSFNGQAECYYFANASEPERCQKMPFNLD 180

PANK3 181 DPYPLLVVNIGSGVSILAVHSKDNYKRVTGTSLGGGTFLGLCSLLTGCESFEEALEMASK 240
Pank3 181 DPYPLLVVNIGSGVSILAVHSKDNYKRVTGTSLGGGTFLGLCSLLTGCESFEEALEMASK 240

PANK3 241 GDSTQADKLVRDIYGGDYERFGLPGWAVASSFGNMIYKEKRESVSKEDLARATLVTITNN 300
Pank3 241 GDSTQADKLVRDIYGGDYERFGLPGWAVASSFGNMIYKEKRESVSKEDLARATLVTITNN 300

PANK3 301 IGSVARMCAVNEKINRVVFVGNFLRVNTLSMKLLAYALDYWSKGQLKALFLEHEGYFGAV 360
Pank3 301 IGSVARMCAVNEKINRVVFVGNFLRVNTLSMKLLAYALDYWSKGQLKALFLEHEGYFGAV 360

PANK3 361 GALLGLPNFS 370
Pank3 361 GALLGLPNFS 370

Supplemental Figure S1

**Homology of PANK3 protein across species.**. Amino acid sequence alignment shows that PANK3 protein is highly conserved between humans and rodents. A. Mouse and human PANK3 are 99.1% homologous, while human and rabbit PANK3 proteins are 100% identical.

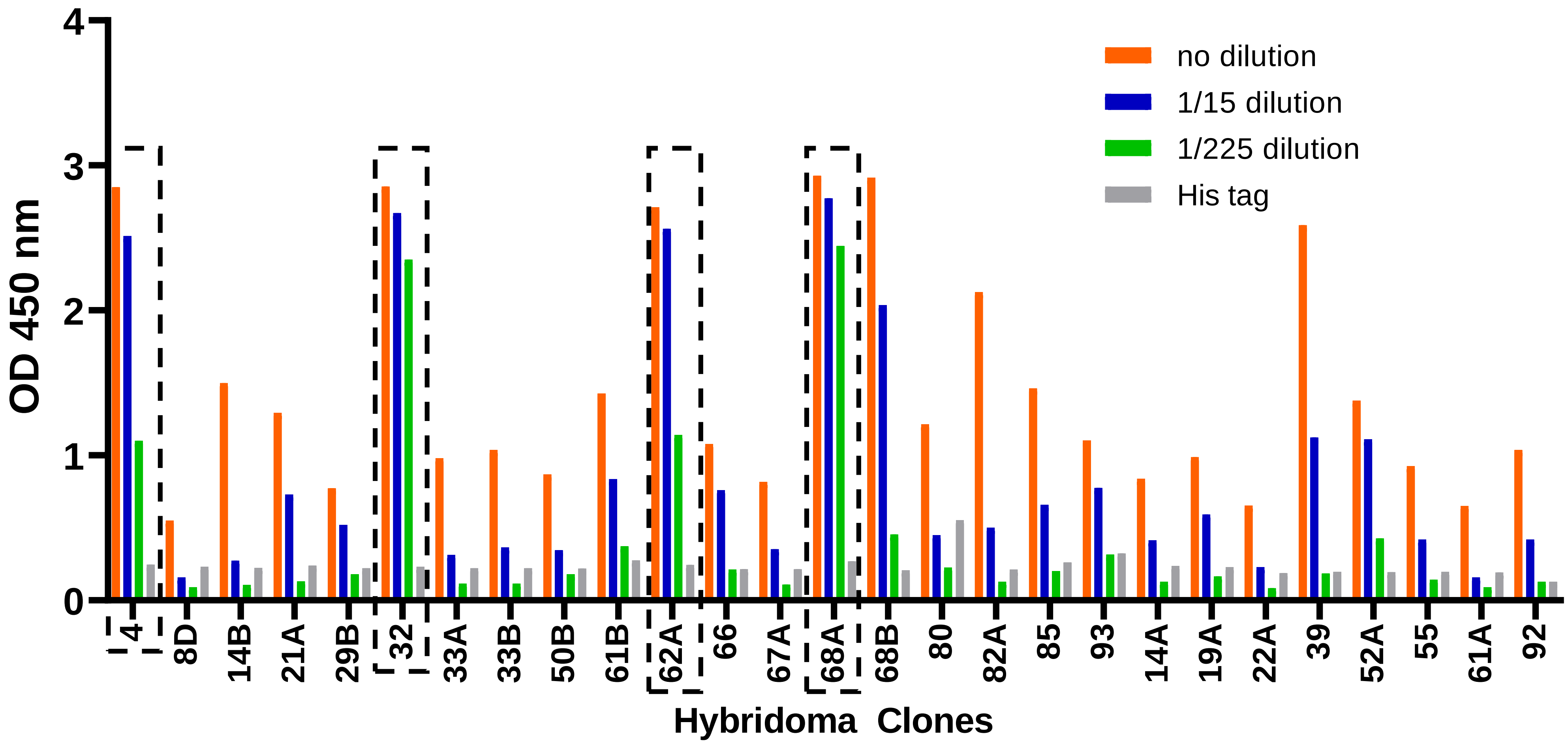

OD 450 nm
0
1
2
3
4
no dilution
1/15 dilution
1/225 dilution
His tag
4
8D
14B
21A
29B
32
33A
33B
50B
61B
62A
66
67A
68A
68B
80
82A
85
93
14A
19A
22A
39
52A
55
61A
92
Hybridoma Clones

Supplemental Figure S2

**Quantification of antibody titre in hybridoma supernatants.** ELISA quantification of media supernatant of all 27 hybridoma clones. Full panel of the hybridoma clones not included in in Figure 4B are shown. The highlighted clones: #4, #32, #62A and #68A were further purified and concentrated based on their ability to detect PANK3 recombinant and endogenous protein in cell lysates by western blots.

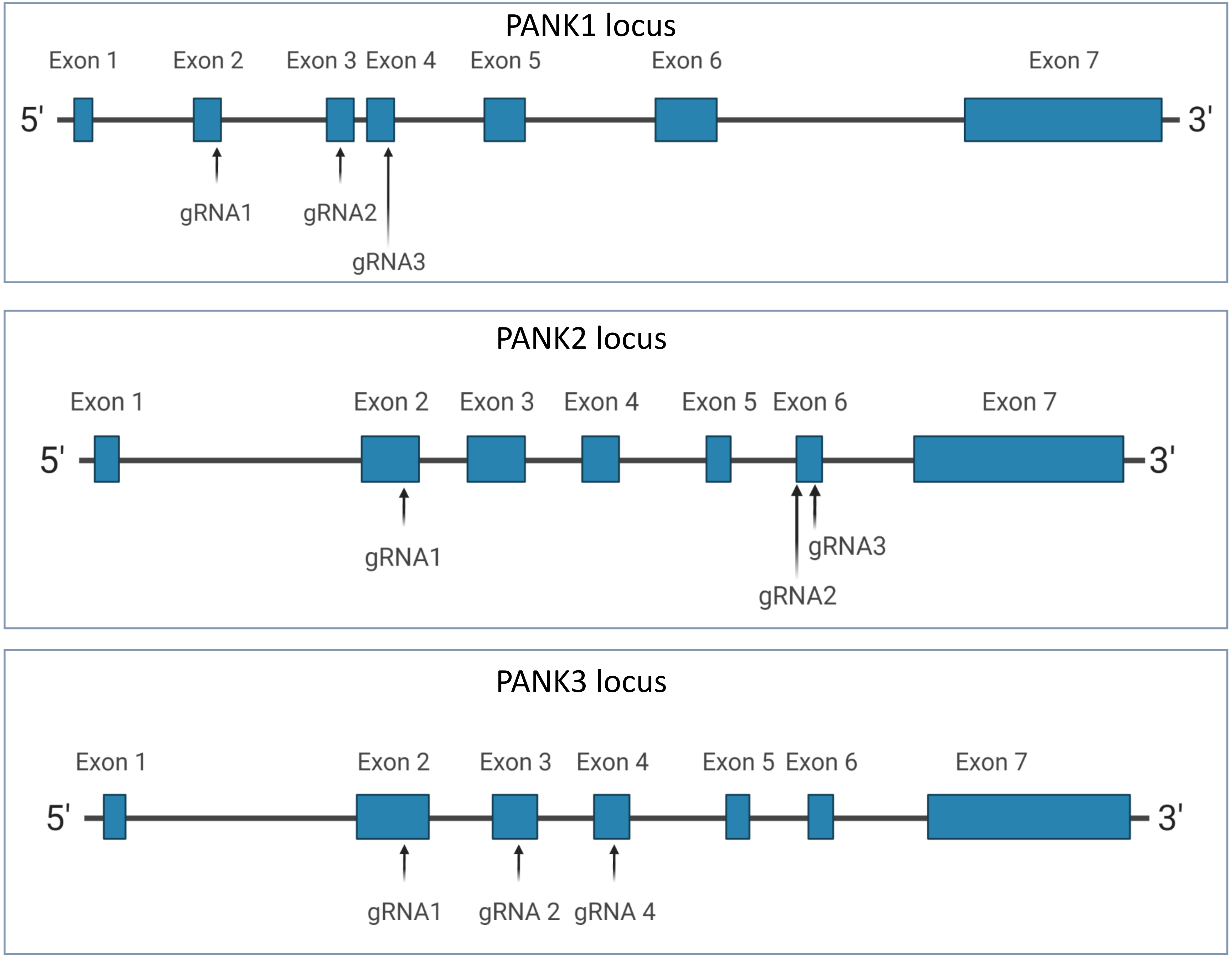
PANK1 locus
Exon 1
Exon 2
Exon 3
Exon 4
Exon 5
Exon 6
Exon 7
5'
3'
gRNA1
gRNA2
gRNA3
PANK2 locus
Exon 1
Exon 2
Exon 3
Exon 4
Exon 5
Exon 6
Exon 7
5'
3'
gRNA1
gRNA3
gRNA2
PANK3 locus
Exon 1
Exon 2
Exon 3
Exon 4
Exon 5
Exon 6
Exon 7
5'
3'
gRNA1
gRNA 2
gRNA 4

Supplemental Figure S3

**PANK1, PANK2 and PANK3 locus identifying 3 different gRNAs that were used to knock the genes out by CRISPR/Cas9 technology.**

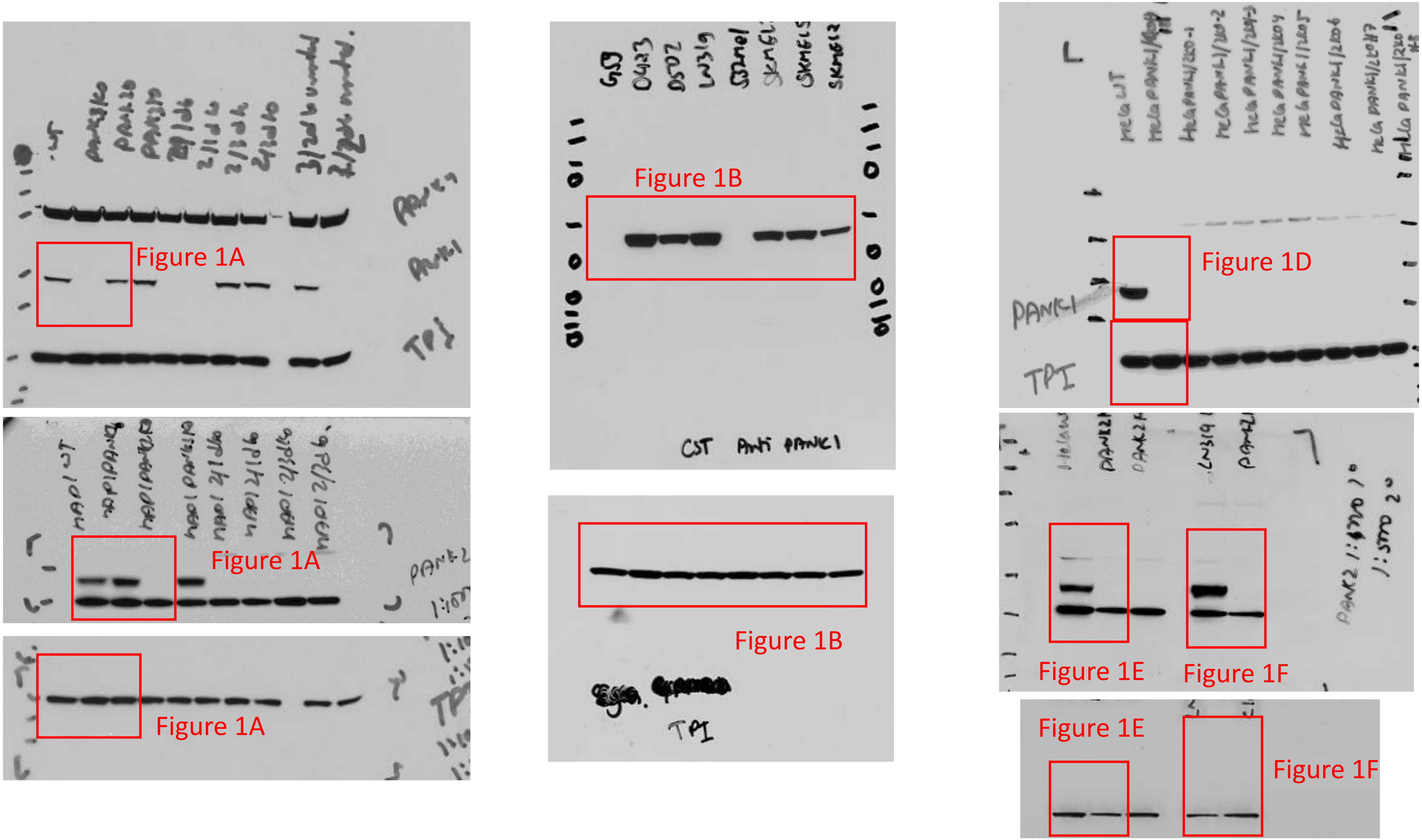

Figure 1A
Figure 1A
Figure 1A
Figure 1B
Figure 1B
Figure 1D
Figure 1E
Figure 1F
Figure 1E
Figure 1F

Supplemental Figure S4

Uncropped pictures of the blots shown in Figure 1. The cropped sections that were used in the figures are highlighted in the red rectangle.

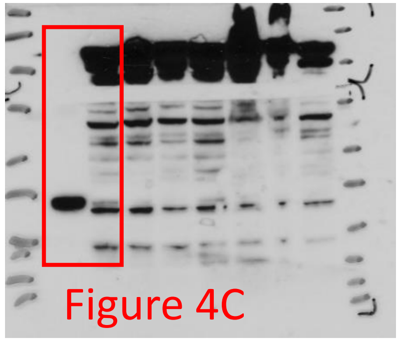
Figure 4C

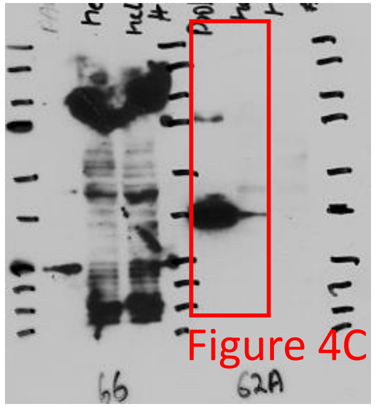
Figure 4C
66
62A

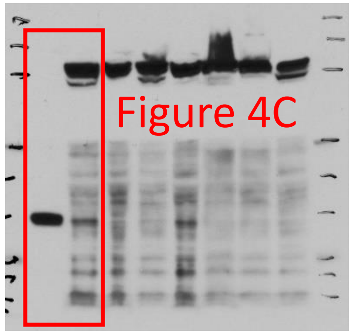
Figure 4C

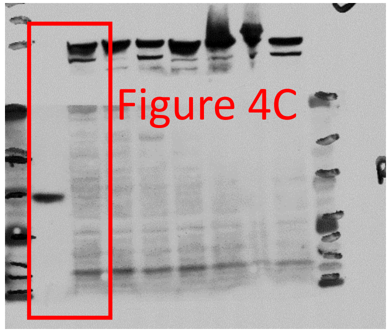
Figure 4C

Supplemental Figure S5

Uncropped pictures of the blots shown in Figure 4. The cropped sections that were used in the figures are highlighted in the red rectangle.

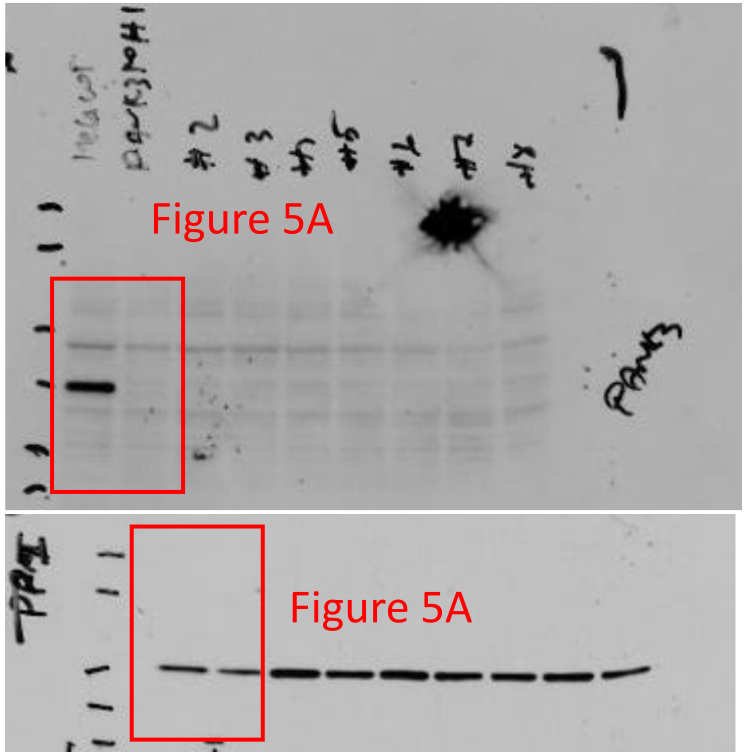
Figure 5A
Figure 5A
PANK3
TPI

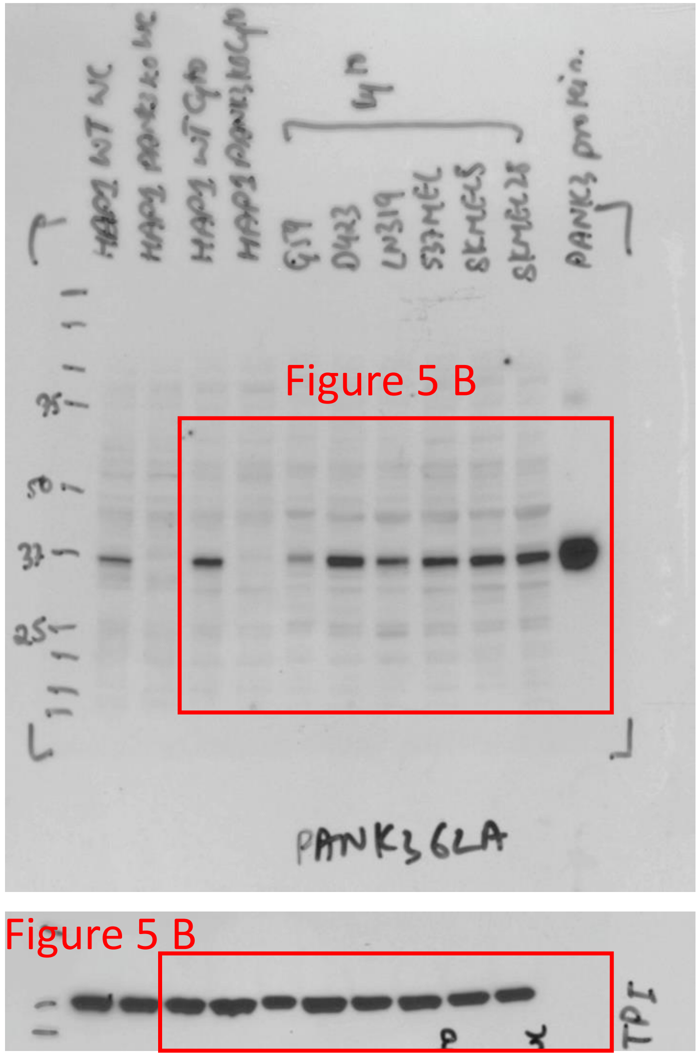
Figure 5 B
Figure 5 B
PANK3 protein
PANK3 6LA
TPI

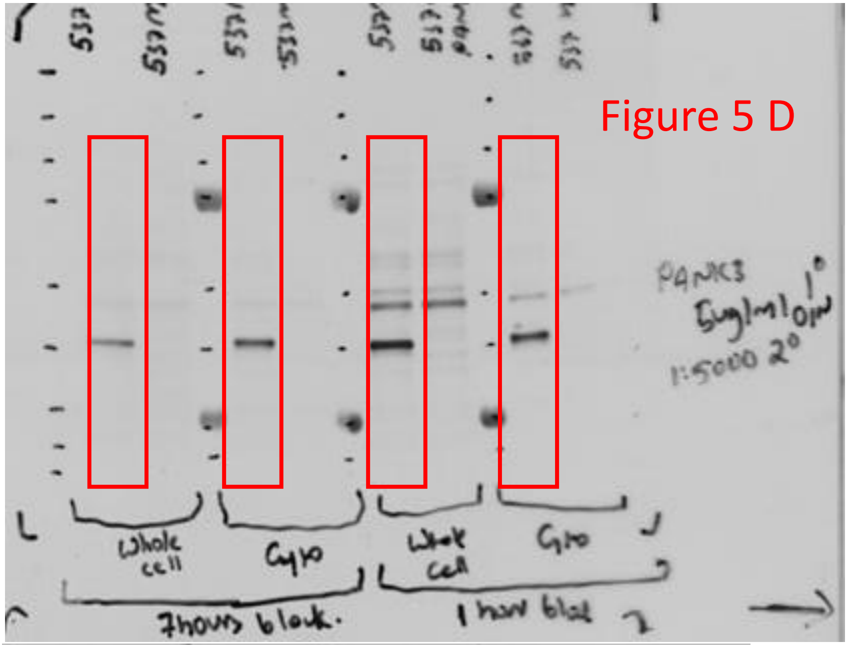
Figure 5 D
Whole cell
Cyto
7hours block
1 hour block
PANK3
1:5000 2°

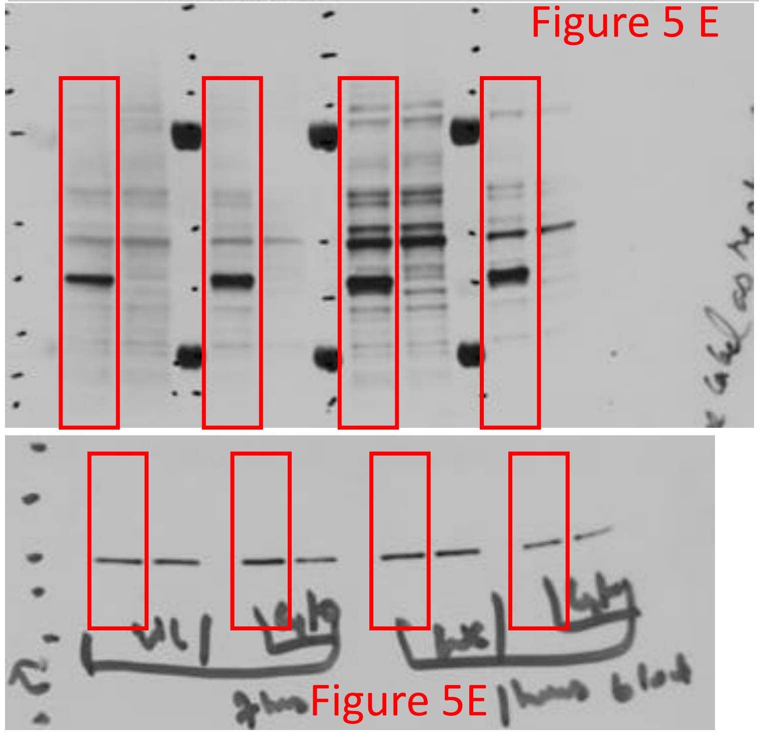
Figure 5 E
Figure 5E

Supplemental Figure S6

Uncropped pictures of the blots shown in Figure 5. The cropped sections that were used in the figure are highlighted in the red rectangle.
